# Phenuivirus glycoprotein Gc domain is a conserved target of cross-functional monoclonal antibodies

**DOI:** 10.64898/2026.10.07.757295

**Authors:** Denise Guerra, Edoardo Cherubini, Mabel R Miranda-Echagüe, Giuseppe Maccari, Gabriele Anichini, Caterina Valoti, Arianna Nurrito, Sandra O Zelech, Emma Ryan, Nicole V Johnson, Gaya K Amarasinghe, Margherita Sambo, Massimiliano Fabbiani, Francesca Montagnani, Maria Grazia Cusi, Jason S McLellan, Ida Paciello, Rino Rappuoli, Emanuele Andreano

## Abstract

Phenuiviruses which include Rift Valley Fever Virus (RVFV), classified by WHO as one of the top priority pathogens with outbreak potential, are emerging arthropod-borne viruses that pose an increasing threat to public health. In line with the “prototype pathogen” framework advanced by the NIAID, the sandfly-transmitted Toscana virus (TOSV) has been adopted as a prototype to advance understanding of relevant phenuiviruses and inform the development of broadly applicable medical countermeasures. In this study, we sought to isolate and characterize monoclonal antibodies (mAbs) elicited by natural TOSV infection to identify shared antigenic determinants. By employing the conserved envelope Gc glycoprotein as bait for antigen-specific B cell sorting, we recovered more than 1,450 memory B cells and identified 112 Gc-specific neutralizing mAbs. Over one-third of these mAbs exhibited cross-reactivity with RVFV, a genetically distinct phenuivirus. Selected mAbs provided prophylactic protection in a murine model of TOSV infection. Together, our findings establish Gc as an important target of protective neutralizing antibodies and provide insights that may guide the rational design of widely effective interventions, strengthening readiness against future phenuivirus outbreaks.

## Introduction

Phenuiviruses, belonging to the *Phenuiviridae* family of the *Bunyavirales* order, are RNA viruses with broad host tropism that infect plants, invertebrates, and humans (Abudurexiti et al., 2019). Human infections are associated with heterogeneous clinical outcomes, from asymptomatic or mild disorders to severe conditions such as thrombocytopenia, hemorrhagic fever and meningoencephalitis (Anywaine et al., 2022; Liu et al., 2014; Nicoletti et al., 1991; Seo et al., 2021). Viral transmission occurs mainly through infected blood-feeding vectors, primarily mosquitoes, ticks and sandflies, although exposure to infected animal blood has also been documented (Ando et al., 2021; Yamanaka et al., 2020). Recently, intensifying globalization and the effects of climate change have facilitated the geographic spread of these arthropods, promoting the emergence of phenuiviruses, such as Rift Valley Fever Virus (RVFV) and Severe Fever with Thrombocytopenia Syndrome Virus (SFTSV), in previously unaffected regions and increasing their public health relevance on a global scale (Azerigyik et al., 2025; Caminade et al., 2018; Elliott, 2009; Mehand et al., 2018; Rolin et al., 2019; Sui et al., 2025).

To support preparedness against these threats, the American National Institute of Allergy and Infectious Diseases (NIAID) has prioritized this viral group under its “prototype pathogen” approach, which focuses on identifying conserved viral features to guide the development of broadly applicable solutions (Cassetti et al., 2023). In this context, Toscana virus (TOSV) has been selected to serve as prototype and inform research on more distantly related phenuiviruses, including RVFV and SFTSV.

First identified in central Italy in 1971 (Verani et al., 1982), TOSV is endemic across the Mediterranean basin, with peak circulation during warmer months (Ayhan et al., 2020; Charrel et al., 2012; Depaquit et al., 2010). Although infection is usually associated with mild symptoms such as headache, nausea and fever (Braito et al., 1997), it can occasionally progress to severe neurological conditions that include meningitis and meningoencephalitis (Ayhan & Charrel, 2020; Nicoletti et al., 1991; Turk et al., 2024). TOSV virions are enveloped particles enclosing a single-stranded RNA genome organized into three fragments, the small, medium and large segments. In addition to non-structural proteins, these segments encode the nucleoprotein, the envelope glycoproteins Gc and Gn, and the RNA-dependent RNA polymerase, respectively (Hornak et al., 2016; Schreur et al., 2018). The Gc and Gn proteins associate as heterodimers that further form pentameric and hexametric arrangements on the viral surface. While Gn is more exposed and sequence-variable, Gc is generally more conserved across related strains (Dessau & Modis, 2013; Halldorsson et al., 2016). These structures mediate receptor binding and membrane fusion during host cell entry and represent major targets for vaccines and antibody-based therapies (Chapman et al., 2023; Gandolfo et al., 2021; Guardado-Calvo & Rey, 2017; Kitandwe et al., 2022; Savellini et al., 2008; Spiegel et al., 2016; Yoshikawa, 2021; Zhao et al., 2025).

Both neutralizing activity and Fc-mediated antibody functions have been implicated in the control and clearance of a broad range of viral infections (DiLillo et al., 2014; Gunn et al., 2018; Lofano et al., 2018; Saphire et al., 2018). Notably, these complementary mechanisms also contribute to humoral immunity against RVFV (Allen et al., 2018; Cartwright et al., 2021; Chapman et al., 2021; Hao et al., 2020; Wang et al., 2019). This highlights the importance of both antibody activity arms and supports their consideration when evaluating protective responses to phenuiviruses, particularly in the context of prototype pathogen studies.

In this study, we characterized the antibody response in subjects with a confirmed TOSV infection to identify shared sites of viral vulnerability that could be translated across phenuiviruses. Using the more conserved TOSV Gc domain as sorting bait, we isolated over 1,450 memory B cells (MBCs) and identified 112 Gc-specific monoclonal antibodies (mAbs) able to neutralize TOSV. Over 35% of these nAbs also showed cross-reactivity against the related RVFV. More than 90% of TOSV Gc-specific mAbs showed Fc-mediated functions and 35% of these antibodies cross-reacted with RVFV. Selected antibody candidates were further assessed *in vivo* using a TOSV infection model, where they conferred protection when administered prophylactically prior to viral challenge. Overall, our data highlights Gc as a conserved target for neutralizing and Fc active antibodies and identifies this domain as a shared site of viral vulnerability that could be leveraged to enhance preparedness against phenuivirus threats.

## Results

### TOSV infection induces an antibody polyclonal response with moderate neutralization and Fc functions

To explore the polyclonal response elicited by TOSV infection, we enrolled four convalescent donors, two males (TOSV_01 and _03) and two females (TOSV_04 and _05), who had a documented TOSV infection between 2022 and 2024. Blood sample collection occurred on day 616, 260, 1,070, and 291 post-infection for subjects TOSV_01, TOSV_03, TOSV_04 and TOSV_05, respectively. Details of the subjects enrolled are provided in **Figure S1**. We initially analyzed the ability of the polyclonal antibody response to bind either the TOSV Gn-Gc complex (**Figure 1A**) or the TOSV Gc antigen alone (**Figure 1B**) using an enzyme linked immunosorbent assay (ELISA). ELISA analysis revealed detectable binding of plasma antibodies from all TOSV convalescent individuals to both recombinant TOSV Gn-Gc and Gc antigens, although marked differences in binding magnitude were observed among donors and between antigens (**Figure 1A-B**). For the Gn-Gc complex, all plasma samples displayed robust, dilution-dependent binding responses, with TOSV_03 showing the highest reactivity, followed by TOSV_05, TOSV_04, and TOSV_01. In contrast, binding to recombinant Gc alone was substantially weaker. While TOSV_03 and TOSV_05 retained high reactivity, TOSV_04 and TOSV_01 exhibited only minimal binding to Gc. Next, to assess the potential cross-reactivity of the polyclonal antibody response, binding to the Gn-Gc glycoprotein complex of RVFV was evaluated by ELISA. None of the plasma samples showed reactivity against the RVFV antigen (**Figure S2A, left panel**), while only the positive control used (RVFV-268) exhibited robust binding (**Figure S2A, right panel**).

**Figure 1.**
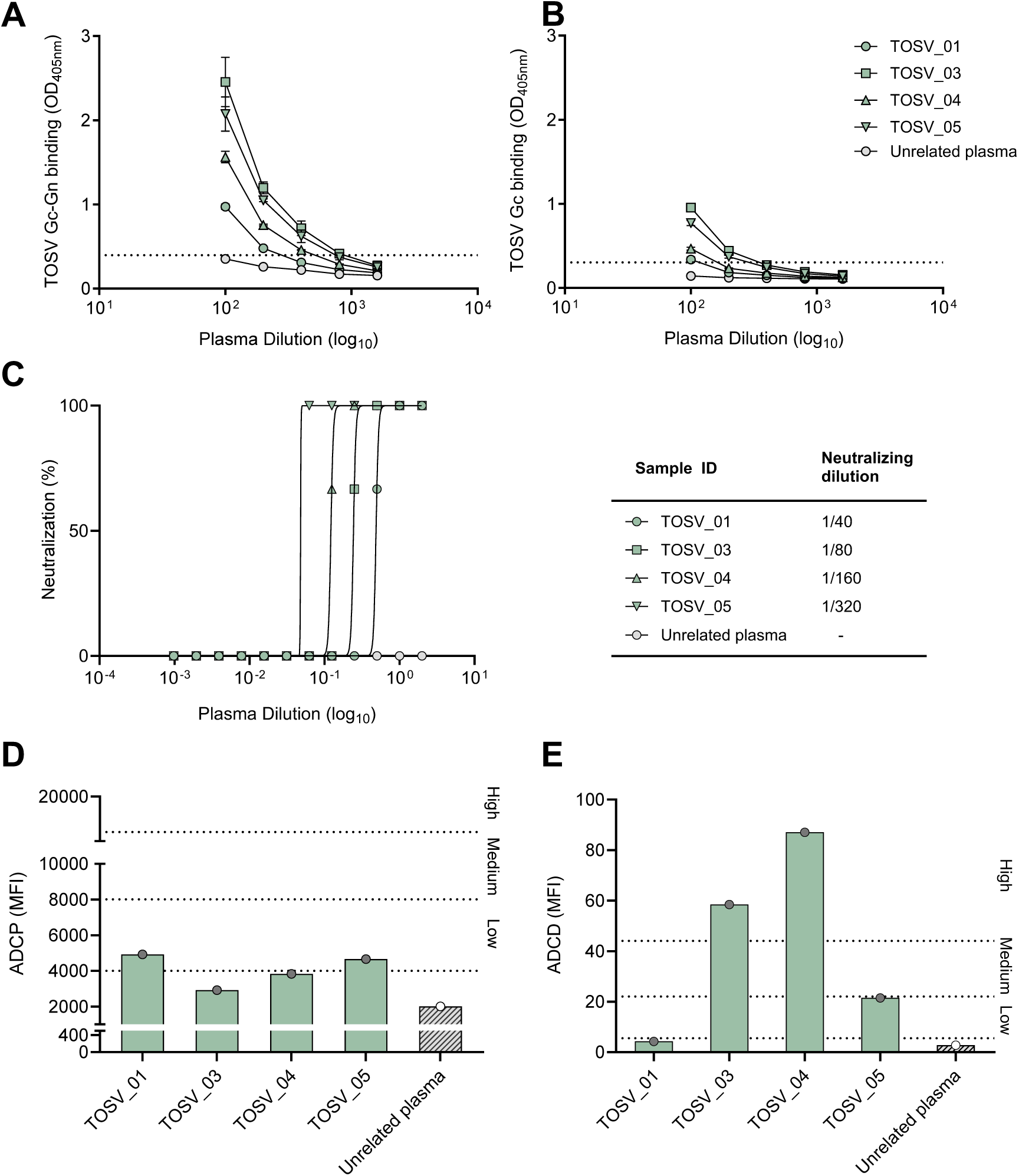
Polyclonal responses in TOSV convalescent donors. (**A-B**) Graphs showing the ability of plasma samples to bind the TOSV Gn-Gc complex (**A**) or the TOSV Gc glycoprotein alone (**B**). Mean and standard deviation are indicated in each graph. Technical replicates were performed for each experiment. Threshold of positivity was set based on an unrelated plasma sample used as negative control. (**C**) Graph showing the neutralizing activity of plasma samples against authentic TOSV (left panel). Technical triplicates were performed for each experiment. Neutralizing dilutions for each subject are reported in the table (right panel). (**D-E**) ADCP and ADCD activity induced by plasma samples derived from TOSV convalescent donors. Unrelated plasma was used as negative control, and threshold of positivity was set accordingly.

The neutralizing activity of plasma samples against authentic TOSV was subsequently evaluated by assessing their ability to protect Vero E6 cell monolayers from virus-induced cytopathic effect. Plasma samples exhibited weak to moderate neutralizing activity, with titers ranging from 1:40 to 1:320. Among all donors, the highest response was observed in subject TOSV_05 (**Figure 1C**).

Considering the relevance of Fc-mediated effector functions to protective immunity against members of the *Phenuiviridae* family (Cartwright et al., 2021), antibody-dependent cellular phagocytosis (ADCP) and antibody-dependent complement deposition (ADCD) assays were performed (**Figure 1D-E**). ADCP activity was assessed by measuring the engulfment of beads coated with fluorescently labeled TOSV Gn-Gc by THP-1 cells. The uptake of antigen-coated beads was confirmed by an increase in the percentage of DY649-positive cells, as determined using the gating strategy shown in **Figure S2B**. Based on the mean fluorescent intensity (MFI) value of an unrelated plasma used as negative control, plasma samples were classified as high (>8-fold the threshold), medium (>4-fold the threshold), or low (>2-fold the threshold) ADCP inducers. ADCP activity was detected only in subjects TOSV_01 and TOSV_05 and remained overall low in magnitude across the cohort (**Figure 1D**). Next, the ability of the polyclonal response to trigger complement activation was quantified by measuring C3b fragment deposition on Gn-Gc-coated beads. As for the phagocytosis assay, ADCD titers were classified as high (>8-fold the threshold), medium (>4-fold the threshold), or low (>2-fold the threshold) based on the unrelated plasma sample used as negative control. The gating strategy applied for data analysis is shown in **Figure S2C**. ADCD activity was observed in 3 out of 4 subjects enrolled in the study. In detail, subject TOSV_04 shows the highest ADCD titers, followed by TOSV_03 and TOSV_05, while plasma from subject TOSV_01 showed no detectable ADCD activity (**Figure 1E**).

### Isolation of 541 Gc-specific MBCs from TOSV convalescent subjects

Peripheral blood mononuclear cells (PBMCs) were collected from all donors and antigen-specific MBC sorting was performed using fluorescently labelled TOSV Gc protein. Starting from live cells, morphological gating and singlet discrimination, CD19⁺ B cells were identified and subsequently gated as CD27⁺IgD⁻, CD27⁺IgD⁻IgM⁻, and CD27⁺IgD⁻IgM⁻Gc⁺ cells (**Figure 2A**). In total, 1,453 Gc-binding MBCs were isolated, with frequencies ranging from 0.13% to 0.27% (**Figure 2B**). Following single-cell sorting, Gc-specific MBCs were cultured on a feeder layer of 3T3 cells expressing CD40L in the presence of IL-2 and IL-21 for 14 days to promote antibody secretion, as previously described (Andreano, Nicastri, et al., 2021; Huang et al., 2013). Then, culture supernatants containing both IgG and IgA antibodies were evaluated for binding to the TOSV Gc subunit by ELISA. This initial screening identified a total of 541 Gc-reactive antibodies covering different signal intensities, indicating substantial variability in antigen recognition (**Figure 2C**). Among the four participants, subject TOSV_03 showed the highest proportion of antigen-specific antibodies (42.2%), followed by TOSV_01 (37.2%), TOSV_04 (33.3%) and TOSV_05 (32.4%), with statistically significant differences observed between donors where indicated.

**Figure 2.**
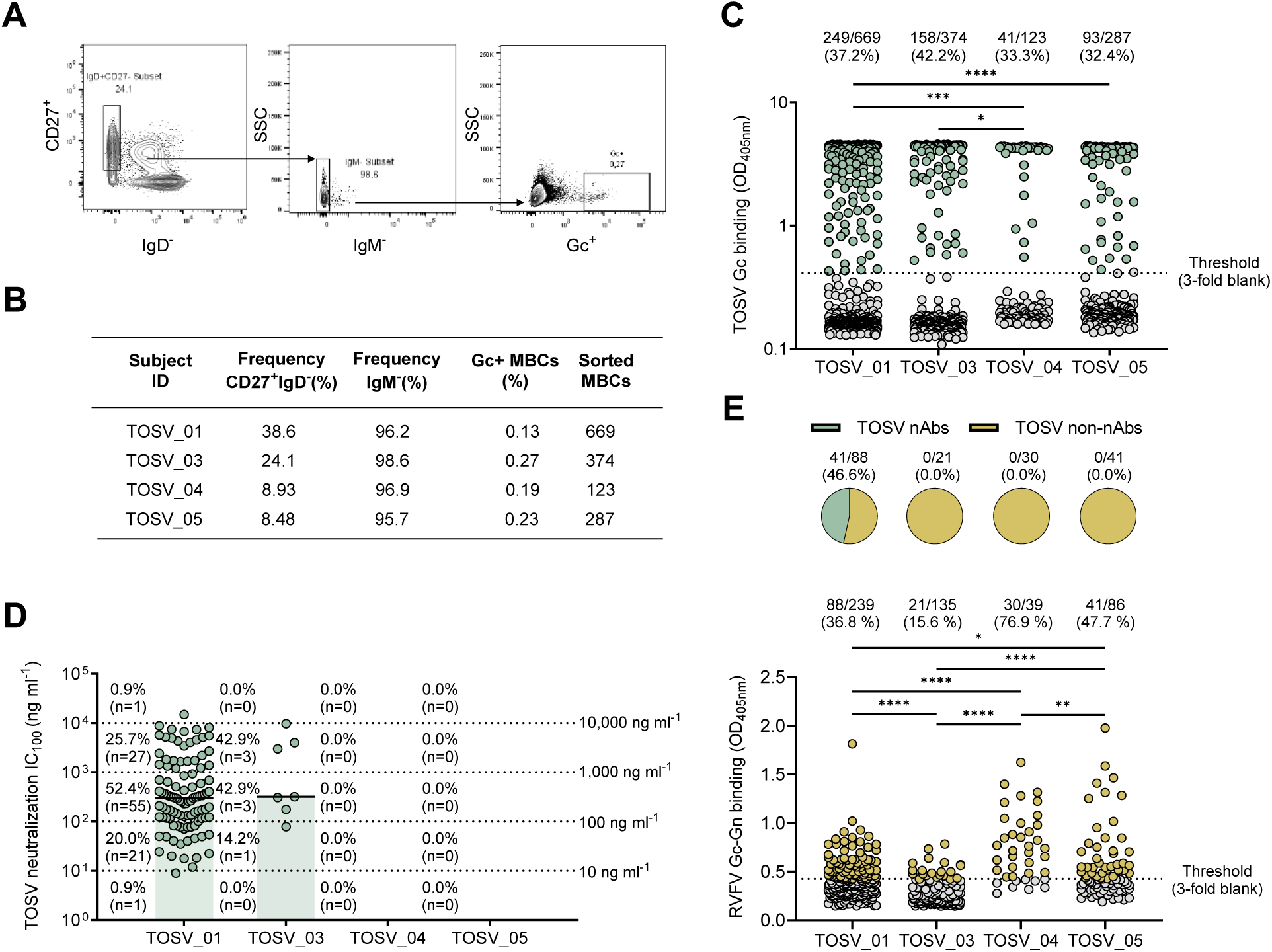
Characterization of antibody responses induced upon TOSV infection. (**A**) The gating strategy used to identify and sort TOSV Gc-specific memory B cells (MBCs) is composed by CD19^+^CD27^+^IgD^-^, CD19^+^CD27^+^IgD^-^IgM^-^ and CD19^+^CD27^+^IgD^-^IgM^-^Gc^+^. (**B**) Frequencies of B cell populations and number of Gc^+^ MBCs sorted from each subject enrolled in this study are reported in the table. (**C**) Binding reactivity of MBC supernatants to the TOSV Gc protein, determined by ELISA and reported as normalized OD_405_ values. The positivity cutoff was defined as three times the signal of the blank control (dotted line). Each point represents one individual supernatant. Green dots indicate Gc-reactive antibodies, whereas grey dots denote non-binding antibodies. A nonparametric Kruskal-Wallis test was used to evaluate the statistical significance between subjects. *P* value significances are shown as \**P* < 0.05, \*\**P* < 0.01, \*\*\**P* < 0.001, and \*\*\*\**P* < 0.0001. (**D**) Scatter plot depicting the neutralizing activity of antibodies against authentic TOSV, expressed as IC_100_ values (ng/mL). Each data point represents one single mAb. All measurements were performed in duplicates. (**E**) Cross-reactivity of expressed mAbs against the RVFV Gn-Gc complex. Threshold of positivity was set based on a negative control (dotted line). Pie charts above the graph show the distribution of RVFV binders that also show neutralizing activity against TOSV. A nonparametric Kruskal-Wallis test was used to evaluate the statistical significance between subjects. *P* value significances are shown as \**P* < 0.05, \*\**P* < 0.01, \*\*\**P* < 0.001, and \*\*\*\**P* < 0.0001.

### TOSV neutralizing mAbs show cross-reactivity to RVFV Gn-Gc

To better understand the neutralization activity and breadth of all TOSV Gc-binding mAbs, we aimed to express all the previously identified 541 antibodies as recombinant IgG1. The immunoglobulin gene expression workflow exhibited an efficiency of 92%, enabling the successful production of 499 mAbs. Expressed Gc-binding mAbs were initially tested in a cytopathic effect-based microneutralization assay (CPE-MN). Each antibody sample was first tested at a single dilution against authentic TOSV, using a viral solution containing 25 median tissue culture infectious dose (25TCID_50_). This analysis identified a total of 112 neutralizing antibodies (nAbs), corresponding to 22.4% of the successfully expressed antibody panel. Notably, the distribution was highly skewed toward subject TOSV_01 (n=105; 93.7%), whereas only a small fraction of nAbs was recovered from donor TOSV_03 (n=7; 6.3%). NAbs were not identified from TOSV_04 and TOSV_05 (**Figure S3**).

Subsequently, all the 112 positive hits were subjected to a secondary CPE-MN assay to investigate their neutralization potency against TOSV using a 100TCID_50_ viral dose. Neutralization potency was quantified based on the concentration required to achieve complete inhibition of viral cytopathic effect (100% inhibitory concentration; IC_100_). Overall, 31 of the identified nAbs (27.7%) displayed limited potency, requiring concentrations above 1,000 ng/mL to achieve full neutralization. A larger subset of antibodies (n=58; 51.8%) exhibited moderate activity, with IC_100_ values ranging from 100 to 1,000 ng/mL, while 22 (19.6%) displayed higher neutralizing capacity, reaching complete viral inhibition at concentrations between 10 and 100 ng/mL. Only one antibody from subject TOSV_01 demonstrated exceptionally high potency, characterized by an IC_100_ value below 10 ng/mL (**Figure 2D**). When tested for binding to the RVFV Gn-Gc glycoprotein complex, 41 neutralizing antibodies showed cross-reactivity to the related phenuivirus (**Figure 2E, top panel**). Overall, considering both nAbs and non-nAbs, 180/499 (36%) of the antibody pool showed reactivity against the RVFV antigen. Among the subjects analyzed, TOSV_04 exhibited a significantly higher cross-reactive response, with 76.9% (30/39) of the isolated mAbs binding to the RVFV Gn-Gc, followed by TOSV_05 (47.7%, 41/86), TOSV_01 (36.8%, 88/239) and TOSV_03 (15.5%, 21/135) (**Figure 2E, bottom panel**).

### TOSV Gc-targeting mAbs induce strong Fc-mediated effector functions

Next, we characterized the ADCP and ADCD activity of the 499 expressed mAbs. Samples were screened as described above and as shown in **Figure 3A**. An unrelated mAb was used as negative control. Antibody titers were classified as high (>64-fold the threshold), medium (>8-to ≤64-fold the threshold), or low (≤8-fold the threshold) based on the positivity threshold established using the unrelated control mAb. Overall, over 90.4% (451/499) of tested mAbs induced detectable phagocytosis against the TOSV Gn-Gc glycoprotein (**Figure 3B, bottom panel**). Notably, antibodies derived from subject TOSV_03 displayed significantly higher ADCP titers compared to those derived from the other donors (**Figure 3B, bottom panel**). Among the antibodies that induced ADCP activity, 44.3% (n=101) from TOSV_01 and 5.3% (n=7) from TOSV_03 were neutralizing (**Figure 3B, top panel**). Moreover, among the ADCP inducers, 83/228 (36.4%), 20/132 (15.1%), 23/31 (74.2%) and 27/60 (45.0%) antibodies from TOSV_01, TOSV_03, TOSV_04 and TOSV_05, respectively, cross-reacted with RVFV (**Figure 3B, middle panel**).

**Figure 3.**
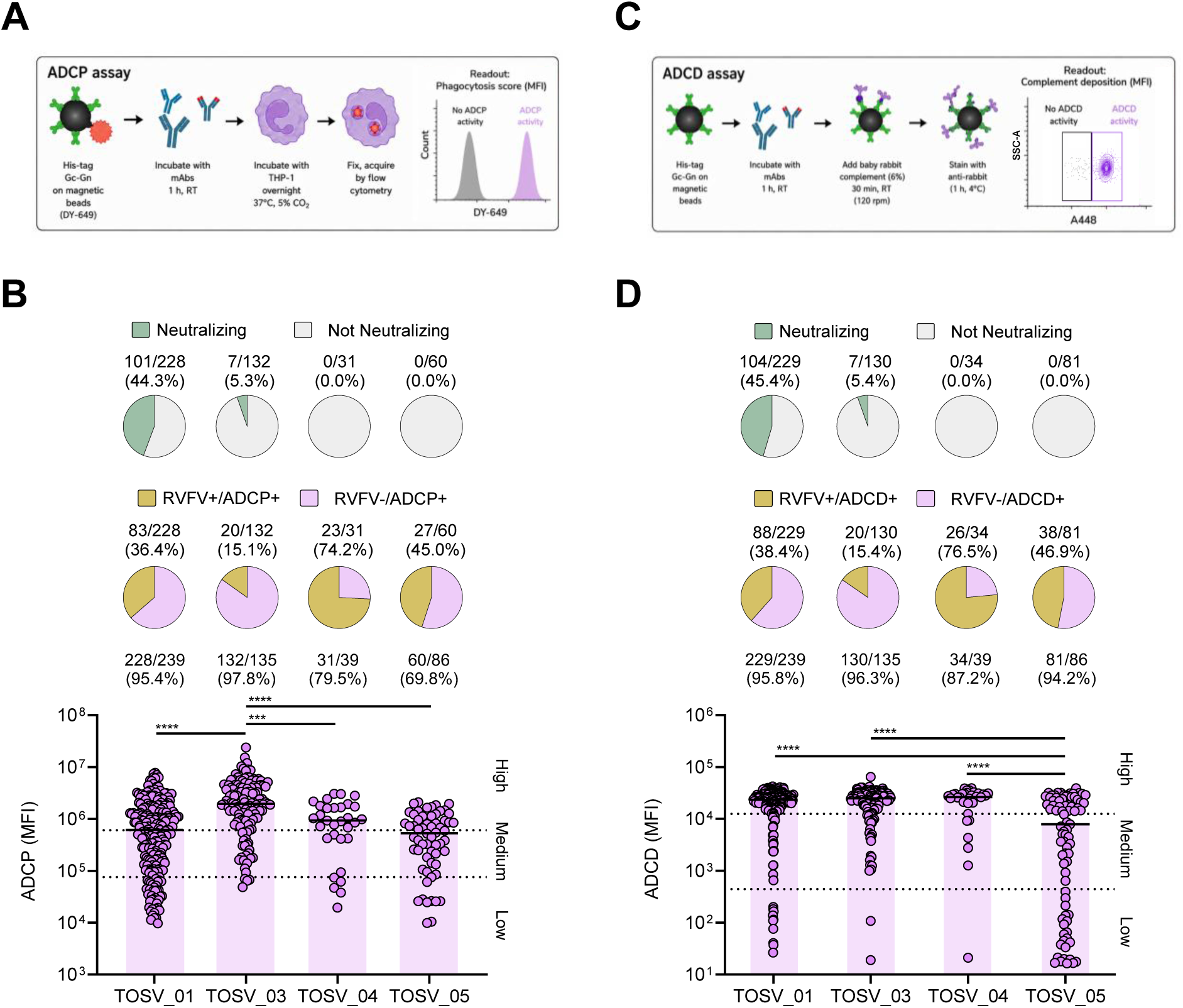
Fc-mediated effector functions of isolated mAbs. (**A**) Schematic representation of the ADCP assay. (**B**) ADCP titers in Gn-Gc mAbs binders. Antibody titers were classified as high, medium, or low based on the positivity threshold established using unrelated mAbs as negative controls. Pie charts in the top panel show the distribution of ADCP-inducing antibodies with neutralizing activity against TOSV. Pie charts in the middle panel show the distribution of ADCP-inducing antibodies with RVFV cross-reactivity. A nonparametric Kruskal-Wallis test was used to evaluate the statistical significance between subjects. *P* value significances are shown as \*\*\**P* < 0.001, and \*\*\*\**P* < 0.0001. (**C**) Experimental workflow used to assess ADCD activity. (**D**) ADCD titers of Gn-Gc mAbs binders. Antibody titers were categorized as high, medium, or low, based on a cutoff defined using unrelated mAbs as negative controls. Pie charts in the top panel depict the proportion of ADCD-inducing antibodies that also exhibit neutralizing activity against TOSV. Pie charts in the middle panel show the distribution of ADCD-inducing antibodies with RVFV cross-reactivity. A nonparametric Kruskal-Wallis test was used to evaluate the statistical significance between subjects. *P* value significances are shown as \*\*\**P* < 0.001, and \*\*\*\**P* < 0.0001.

The same mAbs panel was used to evaluate ADCD following the protocol previously described and outlined in **Figure 3C**. Antibody titers were classified as low (≤8-fold the threshold), medium (>8–28-fold the threshold), or high (>28-fold the threshold), based on the positivity cut-off determined using unrelated mAbs as negative controls. The majority of antibodies isolated from each donor mediated complement deposition, with a frequency of ADCD-positive antibodies comparable across subjects, ranging from 87.2% to 96.3% of the total antibodies tested. Notably, high ADCD activity predominated across all subjects, while intermediate and low responses were observed only in a minority of antibodies (**Figure 3D, bottom panel**). Among the antibodies identified as inducing ADCD activity, 45.4% and 5.4% from subjects TOSV_01 and TOSV_03, respectively, also exhibited neutralizing activity against TOSV (**Figure 3D, top panel**) and 88/229 (38.4%), 20/130 (15.4%), 26/34 (76.5%) and 38/81 (46.9%) from TOSV_01, TOSV_03, TOSV_04 and TOSV_05, respectively, showed binding to RVFV (**Figure 3D, middle panel**).

### Extensive heavy chain mutations of predominant germlines drive TOSV neutralization and RVFV cross-reactivity

The genetic landscape of isolated mAbs was characterized by analyzing the B cell repertoire. A set of 390 heavy chain (HC), 369 light chain (LC) and 290 HC/LC paired sequences were successfully recovered, clustered based on their germline lineages and mapped into a force-directed sequence-similarity network (**Figure 4A**). This topological clustering highlighted the distribution and relative frequencies of neutralizing and RVFV cross-reactive gene usage across the dominant germline lineages. Specifically, neutralizing phenotypes are heavily enriched within distinct germline clusters, such as IGHV4-31 paired with either IGHKV4-1 or IGLV1-40, or IGHV3-23 paired with IGKV1-33. In addition to representing the predominant germlines among TOSV neutralizing antibodies, these B cell lineages were also among the most frequently observed among antibodies cross-reactive with RVFV.

**Figure 4.**
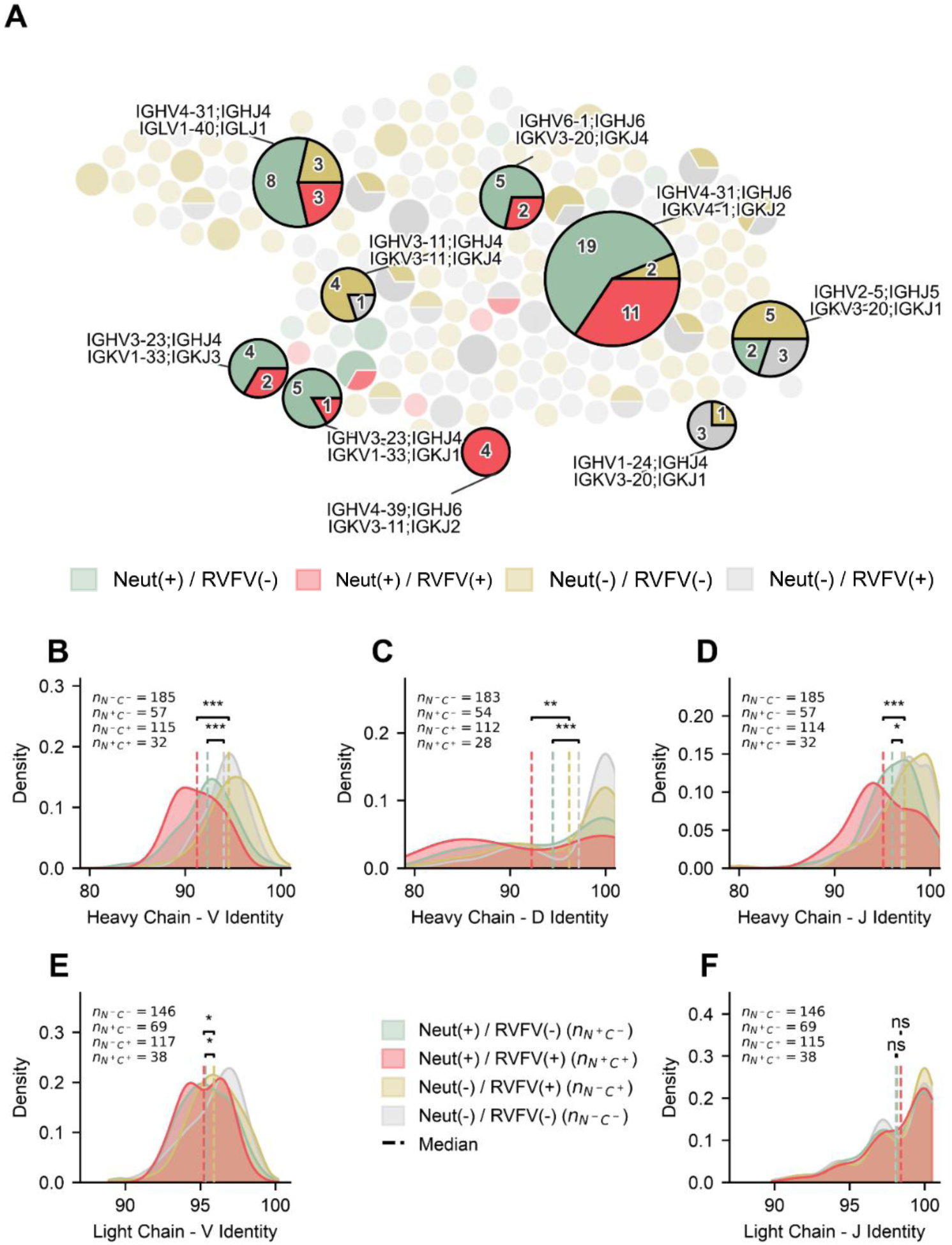
Germline lineage topology and somatic hypermutation profiles of neutralizing and cross-reactive antibodies. (**A**) Force-directed network visualization of heavy and light chain V-J germline pairings. Each circular node represents a distinct germline lineage pairing, with the overall node size proportional to the cluster size. Internal pie charts represents the phenotypic composition within each lineage: RVFV-specific neutralizing (Neut(+)/RVFV(-), green), cross-reactive neutralizing (Neut(+)/RVFV(+), red), cross-reactive non-neutralizing (Neut(-)/RVFV(+), yellow), and double-negative (Neut(-)/RVFV(-), grey). Prominent expanded lineages are annotated with their respective heavy and light chain V and J gene assignments. (**B–F**) Distributions of germline sequence identity percentages for heavy chain V (**B**), D (**C**), and J (**D**) segments, as well as light chain V (**E**) and J (**F**) segments. Vertical dashed lines denote the median germline identity for each respective phenotypic group. Statistical significance between neutralizing and non-neutralizing cohorts was determined using a two-sided Mann-Whitney U test (p < 0.05; *p < 0.01; p < 0.001; ns, not significant).

To determine whether neutralization capacity was associated with specific somatic hypermutation (SHM) profiles, we assessed germline sequence identity across the V, D and J gene segments of the HC and LC (**Figure 4B-F**). As for the HC, neutralizing antibodies exhibited a pronounced and statistically significant increase in mutational burden, compared to their non-neutralizing counterparts. This trend was highly consistent across the heavy V, D and J segments (p < 0.001 to p < 0.05) and was observed independently of antibodies RVFV cross-reactivity. However, nAbs cross-reacting to RVFV Gn-Gc show higher SHM profile compared to nAbs targeting only TOSV. On the other hand, LCs displayed a more conserved mutational landscape. While a minor but significant divergence was detected in the LC V segment for the RVFV(-) background (p < 0.05), LC J identities showed no significant variation between neutralizing and non-neutralizing subsets (**Figure 4E-F**).

### Prophylactic transfer of selected nAbs reduces viral load in TOSV infected mice

To evaluate the protective efficacy of selected nAbs against TOSV challenge, a prophylactic *in vivo* study was conducted with five promising antibody candidates. Neutralizing mAbs were selected based on their potency in the *in vitro* CPE-MN assay and their cross-reactivity with the RVFV Gn-Gc glycoprotein (**Table S1**). Female BALB/c mice received the selected nAbs by intraperitoneal (IP) administration one day prior to subcutaneous TOSV challenge. To monitor viral kinetics following infection, sequential blood samples were collected on day 2 and 5 post-infection (p.i.), as shown in the experimental design (**Figure 5A**). To assess the viral RNA levels in the blood samples, quantitative RT-PCR cycles were performed and expressed as cycle threshold (Ct) values. These were used as a surrogate measure of viral load, with higher Ct values corresponding to lower levels of viral RNA detected in the blood sample.

**Figure 5.**
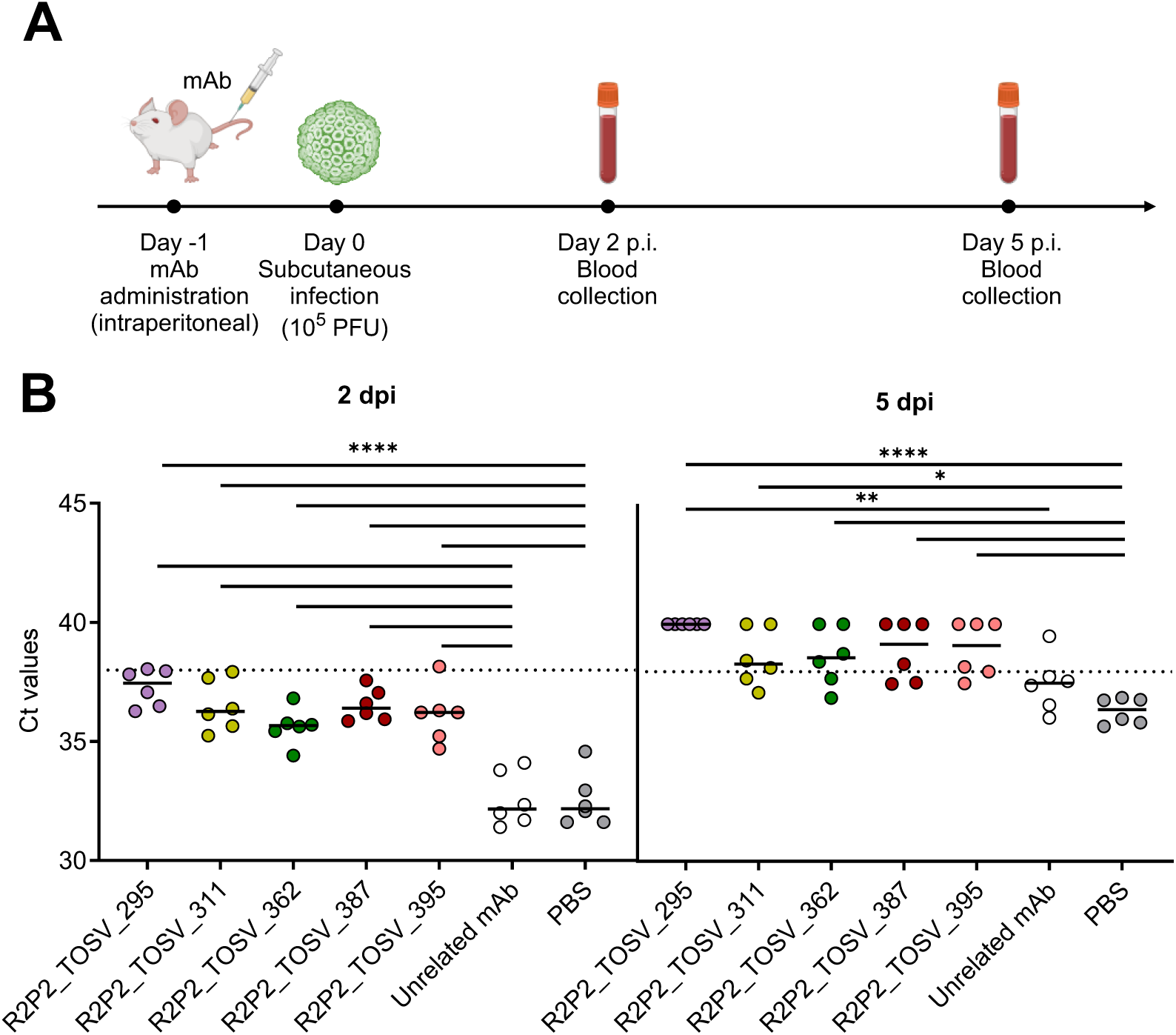
Prophylactic administration of selected nAbs reduces viral load in TOSV-infected mice. (**A**) Schematic representation of the prophylactic in vivo study design. (**B**) Viral RNA levels in individual animals are shown as cycle threshold (Ct) values. The dotted line indicates the assay detection threshold for viral RNA. PBS and an unrelated mAb were included as controls. Each data point represents one animal. Black horizontal lines indicate the mean Ct value for each experimental group. dpi, days post-infection.

Overall, all tested nAbs demonstrated protective activity *in vivo* when administered prophylactically (**Figure 5B**). On both days 2 and 5 p.i., animals treated with TOSV-specific nAbs exhibited higher Ct values compared to the control groups receiving PBS or an unrelated mAb, indicating reduced viral RNA levels in the blood samples. By day 5 p.i., viral RNA was essentially undetectable in most nAb-treated mice, with Ct values exceeding the detection threshold of the assay, indicating robust protection mediated by the administered mAbs. Among the tested mAbs, R2P2_TOSV_295 conferred the strongest protective effect, as evidenced by consistently higher Ct values relative to the other groups. This was in accordance with its high *in vitro* neutralizing potency being the only antibody with an IC_100_ value below 10 ng/mL. Notably, three of the five tested mAbs (R2P2_TOSV_362, 387 and 395) which displayed cross-reactivity against RVFV Gn-Gc, retained protective efficacy in the TOSV challenge model.

## Discussion

In accordance with the NIAID “prototype pathogen” strategy, TOSV has been selected as the representative phenuivirus to inform the rational design of broad interventions against related and more pathogenic members of the same viral family. Insights gained from TOSV research are expected to facilitate the identification of conserved viral vulnerabilities that can be exploited for the accelerated development of vaccines and antibody-based therapeutics with cross-phenuivirus efficacy, with SFTSV representing the primary target for the translation of these efforts. Among the TOSV antigens, the Gc glycoprotein emerges as a particularly attractive target owing to its high degree of conservation across phenuiviruses and its potential to elicit humoral responses with broad recognition capacity (Dessau & Modis, 2013; Halldorsson et al., 2016; Hulswit et al., 2021).

Consistent with this hypothesis, the use of recombinant TOSV Gc as bait for single MBC sorting enabled the identification of more than 540 Gc-specific mAbs, of which 112 displayed neutralizing activity against authentic TOSV. Among the nAb panel, more than 35% also recognized the distantly related RVFV, suggesting the presence of conserved epitopes shared between the two viruses. Furthermore, among the antibodies that retained binding to RVFV Gn-Gc complex, over 36% also induced ADCP and ADCD, highlighting the importance of Fc effector functions as crucial components of humoral immune responses against phenuiviruses. In addition, repertoire analysis revealed that TOSV nAbs are predominantly encoded by different germlines, including IGHV4-31 and IGHV3-23, and that extensive SHM within the HC is a primary driver of the neutralizing phenotype and RVFV cross-reactivity in this cohort, whereas LC maturation plays a secondary role.

When tested in an *in vivo* murine model, both TOSV-specific nAbs and a subset of RVFV cross-reactive mAbs conferred prophylactic protection, indicating that the observed cross-reactivity is not limited to serological recognition but can also mediate functional antiviral protection.

This study has some limitations. First, only four donors were enrolled, and most of the neutralizing antibodies identified were derived from a single individual. Therefore, additional studies involving larger cohorts will be needed to better define the breadth of the antibody response and to characterize the antibody gene rearrangements associated with neutralization and cross-reactivity across different phenuiviruses.

Collectively, the high level of conservation of this glycoprotein across diverse phenuiviruses, together with its capacity to induce cross-functional and protective antibody responses, points to the existence of shared antigenic determinants that are likely maintained by structural and functional constrains, and position Gc as a promising target for the development of broad-spectrum phenuivirus countermeasures.

## Materials and Methods

### Donor enrolment and human sample collection

Human samples from Toscana virus (TOSV) convalescent donors were collected in collaboration with the Azienda Ospedaliera Universitaria Senese (Siena, Italy). The study, initially submitted by Fondazione Toscana Life Sciences, was approved by the local ethics committees (protocol number 24813) and conducted in accordance with Good Clinical Practice and the principles of the Declaration of Helsinki. Written informed consent was obtained from all participants prior to enrolment. The study was non-randomized and unblinded. Peripheral blood mononuclear cells (PBMCs) and plasma were collected from two male and two female TOSV convalescent donors. Plasma samples were heat-inactivated at 56°C for 30 min prior to downstream assays.

### Cells and virus used in this study

Vero E6 cells (African green monkey kidney cell line; ATCC #CRL-1586) were maintained in high-glucose Dulbecco’s Modified Eagle’s Medium (DMEM) (Euroclone, Italy), supplemented with 2 mM L-glutamine (Lonza, Milano, Italy), 100 U/mL penicillin, 100 μg/mL streptomycin (Lonza, Milano, Italy) and 10% fetal bovine serum (FBS) (Euroclone, Italy). Cells were cultured at 37°C, in a humified 5% CO_2_ incubator and routinely passaged every 3-4 days to maintain optimal growth conditions.

For recombinant mAb expression, Expi293F (Thermo Fisher #A14527) and ExpiCHO-S (Gibco #A29127) cells were used. Cells were maintained in their respective manufacturer-recommended culture media and cultured at 37°C, 8% CO₂ with shaking at 125 rpm, according to the manufacturer’s instructions (Thermo Fisher Scientific). Prior to transient transfection, cells were routinely passaged and maintained under optimal growth conditions to ensure high viability and density. Monocytic THP-1 cell line (ATCC #TIB-202) were used for phagocytosis assays and maintained in their manufacturer-recommended culture media.

The TOSV strain 1812, originally isolated from the cerebrospinal fluid (CSF) of a patient with meningitis, was used for this study. The initial isolate underwent two passages in Vero E6 cells (ATCC #CCL-81) and one in McCoy cells (ATCC #CRL-1696) to establish a working stock. To enhance neurovirulence for the murine model, a series of *in vivo* passages was performed. Newborn BALB/c mice (Charles River, Milan, Italy) were inoculated intracerebrally (i.c.) with 10^3^ PFU of the virus. Seven days post-infection, brains were harvested, homogenized on ice using a Dounce homogenizer in Minimum Essential Medium (MEM; Life Technologies, Milan, Italy) supplemented with 2% FBS, and clarified by centrifugation at 2000 g for 20 min at 4°C. This brain homogenate was used for a subsequent i.c. passage in newborn mice, followed by a final passage in adult BALB/c mice. The resulting neuroadapted virus, designated TOSV strain 1812V (Savellini et al., 2008), was propagated in Vero E6 cells to produce a high-titer challenge stock (1.2×10^6^ PFU/mL) and utilized for all other experiments.

### TOSV and RVFV antigen protein expression and purification

The sequence corresponding to Gc residues 831–1281 from TOSV strain SI-1812 (GenBank: OR105998.1) was codon optimized for mammalian expression and subcloned into the mammalian expression vector pcDNA3.1(+) (GenScript). The construct contained an artificial signal sequence (MRPTWAWWLFLVLLLALWAPARG) (Barash et al., 2002) at its N-terminus and an 8 × His tag and Twin-Strep tag, both downstream of a human rhinovirus 3C cleavage site, at its C-terminus. The TOSV Gn-Gc construct was produced by linking codon-optimized TOSV SI-1812 residues 303–610 to Gc residues 831–1281 through a flexible 4 × GGGS/T linker that contained a Strep-II tag between two thrombin cleavage sites. The N-terminus of the Gn-Gc construct contained the same artificial signal sequence as the Gc construct, whereas the C-terminus contained an 8 × His tag.

TOSV proteins were expressed by GenScript in the TurboCHO™ platform and purified from crude supernatant using the HisTrap™ FF Crude resin (Cytiva). The purity and homogeneity of the antigens were assessed by reducing and non-reducing SDS-PAGE and SEC-HPLC with a TSKgel G3000SWxl column (Tosoh Bioscience) using 0.1 M Na2SO4 in 0.12 M phosphate buffer (pH ∼6.7) as the mobile phase. Purified antigens were buffer-exchanged into PBS (pH 7.2) before storage at -70 °C.

RVFV Gn-Gc (aa 154-469 and 691-1114; strain ZH501, accession DQ380200) was engineered as a single-chain ectodomain construct linked by a 44-residue flexible linker containing thrombin cleavage sites and an internal Strep tag, as previously described (Fatma et al., 2026). Recombinant Gn-Gc proteins were expressed in Expi293F (GnTI-) cells and purified by Ni-affinity, anion-exchange, and size-exclusion chromatography. Protein purity, integrity, and identity were confirmed by SDS-PAGE, Coomassie Blue staining, and mass spectrometry.

### Peripheral blood mononuclear cell isolation and memory B cell sorting

PBMCs were isolated as previously described (Andreano, Nicastri, et al., 2021). Briefly, PBMCs were obtained from heparinized whole blood by density-gradient centrifugation using Ficoll-Paque PREMIUM (GE Healthcare). Then, cells were incubated at room temperature (RT) with Live/Dead Fixable Aqua (Invitrogen) diluted 1:500 in phosphate-buffered saline (PBS; Gibco). After 20 min, PBMCs were washed with PBS, and nonspecific binding was blocked with 20% normal rabbit serum (Life Technologies) for 30 min at 4°C. To identify Gc⁺ MBCs, PBMCs were stained for 20 min at 4°C with the TOSV Gc antigen labelled with Strep-Tactin^®^XT DY-488 (IBA Lifesciences). Cells were additionally stained at 4°C with the following antibodies: CD19 BV421, IgM PerCP-Cy5.5, CD27 PE and IgD Alexa Fluor 700 (BD Biosciences), CD3 PE-Cy7, CD14 PE-Cy7 and CD56 PE-Cy7 (BioLegend). After washing and resuspension in sorting buffer (PBS supplemented with 2.5 mM EDTA), MBCs (CD3⁻CD14⁻CD56⁻CD19⁺CD27⁺IgD⁻IgM⁻) were single-cell sorted using a BD FACSAria Fusion (BD Biosciences) into 384-well plates containing 3T3-CD40L feeder cells. Sorted cells were cultured for 14 days in the presence of IL-2 and IL-21, as previously described (Andreano, Nicastri, et al., 2021; Huang et al., 2013).

### ELISA assay of sorting supernatants, plasma samples and recombinant mAbs

The presence of TOSV Gc-binding antibodies in culture supernatants derived from sorted Gc^+^ MBCs was evaluated by an initial ELISA assay. Briefly, high-binding 384-well flat-bottom microtiter plates (Greiner Bio-one) were incubated overnight at RT with 25 μL/well of streptavidin (Thermo Fisher Scientific) diluted in PBS to a final concentration of 3 μg/mL. The following day, plates were washed three times with washing buffer (1X PBS + 0.05% Tween20), and subsequently coated with 25 μL/well of TOSV Gc antigen at 3 μg/mL in PBS for 1h at RT. After three additional washes, 50 μL/well of saturation buffer (1X PBS + 1% BSA) were added to the plates to minimize non-specific binding, and plates were incubated for 1h at 37°C without CO_2_. For this initial screening, 5 μL of culture supernatants were diluted in 20 μL of dilution buffer (1X PBS + 1% BSA + 0.05% Tween20), added to the wells and incubated for 1h at 37°C without CO_2_. Following incubation, plates were washed three times and incubated with 25 μL/well of secondary goat anti-human IgG (Sigma-Aldrich) and IgA (Southern Biotech) conjugated with alkaline phosphatase (diluted 1:2000 in dilution buffer) for 1h at 37°C without CO_2_. After three final washing steps, 25 μL/well of p-nitrophenyl phosphate substrate (pNPP) (Sigma-Aldrich) were added to detect TOSV Gc-binding antibodies, and the final reaction was quantified by measuring absorbance at a wavelength of 405 nm using a Varioskan Lux Microplate Reader (Thermo Fisher Scientific). Plasma from TOSV convalescent donors served as positive controls, while unrelated plasma and a SARS-CoV-2-specific mAb were used as negative controls. The threshold for sample positivity was defined as an optical density value at least threefold higher than that of the blank.

Similarly, ELISA assays were performed to assess binding of plasma samples from the four TOSV convalescent subjects and recombinant mAbs against the TOSV Gn-Gc complex, the TOSV Gc alone and the RVFV Gn-Gc glycoproteins. Plasma samples from TOSV convalescent donors were tested starting at a 1:100 dilution followed by two-fold serial dilutions, whereas recombinant mAbs were tested at a 1:5 dilution. For screenings involving the RVFV antigen, the RVFV-268 mAb (Chapman et al., 2021) was included as a positive control, and an unrelated plasma served as a negative control in all experiments. All samples were tested in duplicate, with sample buffer used as a blank. The threshold for positivity was defined as an optical density (OD) value at least three-fold higher than that of the blank.

### Single-cell RT-PCR, immunoglobulin gene amplification, and transcriptionally active PCR (TAP)-mediated expression of mAbs

Single-cell reverse transcription polymerase chain reaction (RT-PCR), immunoglobulin gene amplification, and transcriptionally active PCR (TAP) expression were undertaken to enable the production of full-length antibodies. Specifically, 5 μL of cell lysate derived from the original 384-well sorting plate were subjected to RT-PCR, followed by two sequential amplification steps (PCRI and nested PCRII), as previously reported (Andreano, Nicastri, et al., 2021; Andreano, Paciello, et al., 2021). The resulting PCRII amplicons were subsequently utilized for the identification of antibody heavy chain (HC) and light chain (LC) sequences via Sanger sequencing, as well as for their cloning into appropriate expression vectors, in accordance with established methodologies (Liang et al., 2002; Tiller et al., 2008). TAP was performed using a reaction mixture comprising 0.25 μL of Q5 polymerase (NEB), 5 μL of GC enhancer (NEB), 5 μL of 5X reaction buffer, 0.5 μL of dNTPs, 0.125 μL of forward and reverse primers, and 3 μL of ligation product. Thermal cycling conditions included an initial denaturation step at 98°C for 2 min, followed by 35 cycles of 98°C for 10 s, 61°C for 20 s, and 72°C for 90 s, with a final extension at 72°C for 5 min. TAP products were purified under conditions analogous to those employed for PCRII, quantified, and subsequently used for transient transfection of Expi293F cells, in strict accordance with the manufacturer’s instructions.

### Authentic TOSV propagation and titration

For viral propagation, Vero E6 cells were seeded one day prior to infection at a density of 1.8×10^5^ cells/mL in a final volume of 50 mL of culture medium in T175 flasks and incubated overnight at 37°C with 5% CO_2_. On the day of the infection, culture medium was removed and the Vero E6 cell monolayer was washed twice with sterile PBS. Cells were then infected with 5 mL of authentic TOSV virus inoculum diluted 1:100 in DMEM supplemented with 2% FBS and incubated for 1 h at 37°C with 5% CO_2_ to allow viral adsorption. At the end of the incubation step, 45 mL of DMEM + 2% FBS were added to the flask. Infected cell cultures were maintained at 37°C with 5% CO_2_ and monitored daily for cytopathic effect (CPE). When CPE reached approximately 80-90%, supernatants were harvested and clarified by centrifugation at 4°C (1600 rpm for 8 min) to remove cell debris. The clarified viral stock was aliquoted and stored at -80°C until further use.

Viral titer was determined by 50% tissue culture infectious dose (TCID_50_) assay using confluent Vero E6 cell monolayers seeded in 96-well plates the day before the titration. Cells were infected with serial 1:10 dilutions of the virus and incubated at 37°C, in a humidified atmosphere with 5% CO_2_. Plates were monitored daily for up to 4 days using an inverted light microscope to assess the development of TOSV-induced CPE. Virus titer was calculated using the Spearman-Karber method (ref) and expressed as the reciprocal of the highest dilution producing CPE in at least 50% of the inoculated wells.

### Authentic TOSV neutralization assay

Neutralization activity of both sorting and TAP-expressed supernatants was initially screened using a qualitative CPE-MN assay at a single point dilution to identify positive candidate samples. In brief, Vero E6 cells were seeded one day prior to infection at a density of 1.5×10^5^ cells/mL in 96-well flat-bottom plates (100 μL/well) in DMEM supplemented with 2% FBS and 1% penicillin/streptomycin and incubated overnight at 37°C with 5% CO_2_. Supernatants were diluted 1:4 in culture medium and mixed with a TOSV viral solution at a final dose of 25TCID_50_. Following a 1h incubation at 37°C with 5% CO_2_ to allow antibody-mediated neutralization, each mixture was transferred onto sub-confluent Vero E6 cell monolayers prepared the previous day. Plates were then incubated for 4 days under standard culture conditions and examined for CPE by visual inspection using an inverted light microscope. The presence or absence of CPE was assessed relative to control wells consisting of infected cells without antibody (positive control) and uninfected cells (negative control).

TAP-expressed samples that demonstrated protective activity in the initial single-point screening were further characterized for their neutralizing potency against authentic TOSV. The assay was conducted as described above, with the exception that the viral input was increased to 100TCID_50_. Samples were tested starting from a 1:4 dilution, followed by two-fold serial dilutions. All conditions were assessed in technical duplicates. The same protocol was used to test neutralization activity of the plasma samples from the four TOSV convalescent subjects. All samples were initially diluted at 1:10, followed by two-fold serial dilutions. Three technical replicates were performed for each plasma sample, and unrelated plasma was used as negative control.

### Antibody-dependent cellular phagocytosis assay (ADCP)

Antibody-dependent cellular phagocytosis (ADCP) was assessed using a flow cytometry-based assay as reported previously (Paciello et al., 2024) with variation in the coating antigen. Briefly, 50 μg of histidine tagged TOSV Gn-Gc protein was labeled using Strep-Tactin™XT Conjugate DY649 and coated with His-tag magnetic beads (Dynabeads His-Tag, Invitrogen) according to the manufacturer’s instructions. The Gn-Gc glycoprotein beads were incubated with 20 µL of mAbs or heat inactivated plasma diluted 1/10 in complete RPMI (RPMI supplemented with 10% heat-inactivated FBS, 1% Penicillin Streptomycin, 1% HEPES and 1% Pyruvate) for 1h at RT. Following, this mix was incubated with the monocytic THP-1 cell line overnight at 37°C with 5% CO_2_. Then, cells were fixed using fixation buffer (BioLegend) according to the manufacturer’s protocol and acquired on BD CANTO (Becton Dickinson). Single technical replicates were performed for mAbs experiments, and two replicates for plasma sample experiments. Unrelated plasma and unrelated mAbs were used as negative control. Phagocytosis score was calculated as the percentage of THP-1 cells that engulfed fluorescent beads multiplied by the median fluorescence intensity (MFI) of the population.

### Antibody-dependent complement deposition assay (ADCD)

Flow cytometry-based assay was performed to measure ADCD. Briefly, 50 µg of histidine tagged TOSV Gn-Gc were coated with 30 µL of His-Tag beads (Dynabeads His-Tag, Invitrogen) as manufacture’s protocol. Then, heat-inactivated plasma and mAbs, diluted 1:10 in PBS (Gibco) were incubated with TOSV Gn-Gc coated beads for 1h at RT. Then, 6% of baby rabbit complement (Cederlane) was added and samples were incubated at RT and 120 rpm for 30 min. After incubation, the immune complex was washed with PBS (Gibco) and stained with goat anti-rabbit polyclonal antibody against C3 (MP Biomedicals) for 1h at 4°C. After washing with PBS, samples were acquired using the BD CANTO (Becton Dickinson). Unrelated plasma and mAbs were used as negative control. Single technical replicates were performed for each experiment. Results were reported as median fluorescence intensity of C3 deposition on the beads.

### Recombinant expression of selected TOSV-specific mAbs for *in vivo* studies

Plasmids encoding the selected mAb HC and LC were transiently co-transfected into 100 mL of ExpiCHO-S™ cells in a 2:3 ratio, following ExpiCHO™ Expression System protocol. Eight days post-transfection, cell cultures were harvested by centrifugation, and supernatants were filtered through a 0.22 μm filter. Antibodies were then purified through affinity chromatography using ÄKTA go chromatography system (Cytiva). Clarified supernatants were loaded in 5 mL HiTrap Protein G HP column (Cytiva) pre-equilibrated with Buffer A (0.02 M NaH2PO4, pH 7). Antibodies were eluted by applying 5 column volumes of Buffer B (0.1 M glycine-HCl, pH 2.7). Eluted fractions (1 mL each) were collected and neutralized with 100 μL of 1M Tris-HCl, pH 9.0. Non-reducing sodium dodecyl sulfate-polyacrylamide gel electrophoresis (SDS-PAGE) was performed to analyze the fractions and identify those containing the antibodies, that were consequently pooled and dialyzed overnight against PBS at 4°C. Final antibody concentrations were determined by measuring protein content using the Pierce BCA Protein Assay Kit (Thermo Fisher Scientific). Purified proteins were stored at −80°C before use.

### Prophylactic mAb transfer to BALB/c mice

Four-week-old female BALB/c mice (Charles River, Milan, Italy) were utilized for all prophylactic experiments, with groups comprising six animals each. The core experimental design involved an initial intraperitoneal (IP) injection of the candidate mAbs (50 mg/kg), with dosages calculated based on the mice’s pre-recorded body weights. Twenty-four hours post mAb injection, animals were challenged subcutaneously with 10^6^ plaque-forming units (PFU) of TOSV. To monitor viral kinetics, serial blood samples were collected via submandibular vein puncture at 2 and 5 days post-infection (p.i.).

### RNA extraction and viral quantification

Viral RNA was extracted from inactivated whole blood and spleen samples using the automated EZ1 DSP Virus Kit (Qiagen) according to the manufacturer’s instructions. TOSV RNA levels were quantified via one-step real-time RT-PCR targeting the nucleoprotein (N) gene using the following sequences: Forward 5’-TGGCTGCTACGTTTGAAAAGC-3’, Reverse 5’-GCTTCTTTTTGTCCTCCCTAGTGA-3’, and Probe 5’-[FAM]-CATGGCTGCCATGAGTGGACGGT-[TAMRA]-3’. To ensure extraction quality and normalization for PCR inhibitors, the murine GAPDH gene was amplified as an internal housekeeping control. Viral loads were calculated as cycle threshold (Ct) values.

### Sequence assembly and annotation

Individual antibody sequence assembly and annotation were performed using a combination of tools and scripts to ensure high quality of the analysis. An initial set of 942 raw paired-end Sanger sequence chromatograms were converted to FASTQ format, and reads were trimmed at both ends using a Phred quality score of 20 as a threshold. Forward and reverse reads were then assembled into contiguous sequences using the AssemblePairs.py module from the pRESTO toolkit (Heiden et al., 2014). During assembly, reverse reads were reverse-complemented, and pairs were merged via an alignment-based consensus approach with a maximum allowed error rate of 0.4. As a result, a total of 781 heavy and light chain sequences were successfully assembled. Assembled Ig sequences were annotated with IgBLAST (Ye et al., 2013) against the human IMGT reference germline databases for V, D and J genes (Giudicelli et al., 2005). The annotated sequences were further subjected to filtering to eliminate incomplete or artifact entries. A sequence was classified as a complete, fully resolved fragment if the following criteria were met: i) unambiguous assignment of V and J genes; ii) presence of a defined CDR3; iii) complete identification of framework regions 1, 2, and 3; presence of an intact FR4, or a J-gene alignment spanning greater than 20 nucleotides. Following this quality control, a final dataset of 699 Ig sequences was retained for downstream analysis.

### Repertoire analyses of TOSV nAbs

Annotated heavy and light chain sequences were parsed and clustered by their respective V-J germline arrangements. To visualize the topological landscape of germline usage, representative CDR3 of full-length sequence were extracted for each cluster and subjected to global pairwise alignment using the BLOSUM62 substitution matrix. Alignment scores were then converted to pairwise distances and transformed into an adjacency matrix using an exponential decay function. A k-nearest neighbor graph with k <5 was conducted in NetworkX (Hagberg et al., 2008) using a force-directed spring layout. Clusters were represented as pie chart to reflect the distribution of neutralization and cross-reactivity. Furthermore, the distribution of V, D and J germline sequence identities were plotted for all four groups (Neutralizing and cross-reactive, non-neutralizing and cross-reactive, neutralizing and non-cross-reactive, non-neutralizing and non-cross-reactive). Statistical differences in median germline identities between cohorts were evaluated using two-sided Mann-Whitney U test, with significance threshold denoted as follow: *(p < 0.05); **(p < 0.01); ***(p < 0.001).

## Acknowledgments

We thank the TOSV convalescent donors for their participation in the study and the staff of the Azienda Ospedaliera Universitaria Senese, Siena, Italy, for collecting and providing the blood samples under study protocols approved by the local ethics committees. This work was financially supported by a grant from the US NIH: U19AI181984.

## Author contributions

Conceptualization: R.R. and E.A.

Methodology: D.G., E.C., M.R.M.E., G.M., G.A., C.V., A.N., S.O.Z., I.P.

Investigation: D.G., E.C., M.R.M.E., G.M., G.A., C.V., A.N., S.O.Z., I.P.

Visualization and data curation: D.G., I.P., E.C., M.R.M.E., G.M., G.A., M.G.C., E.A.

Resources: E.R., N.V.J., G.K.A, M.S., M.F., F.M., J.S.M.

Funding acquisition: M.G.C., R.R., E.A.

Supervised the work: M.G.C., J.S.M., I.P., R.R., E.A.

Writing – original draft: D.G., I.P., E.C., M.R.M.E., G.M., E.A.

Writing – review and editing: all authors.

## Supplementary Figures

**Figure S1.**
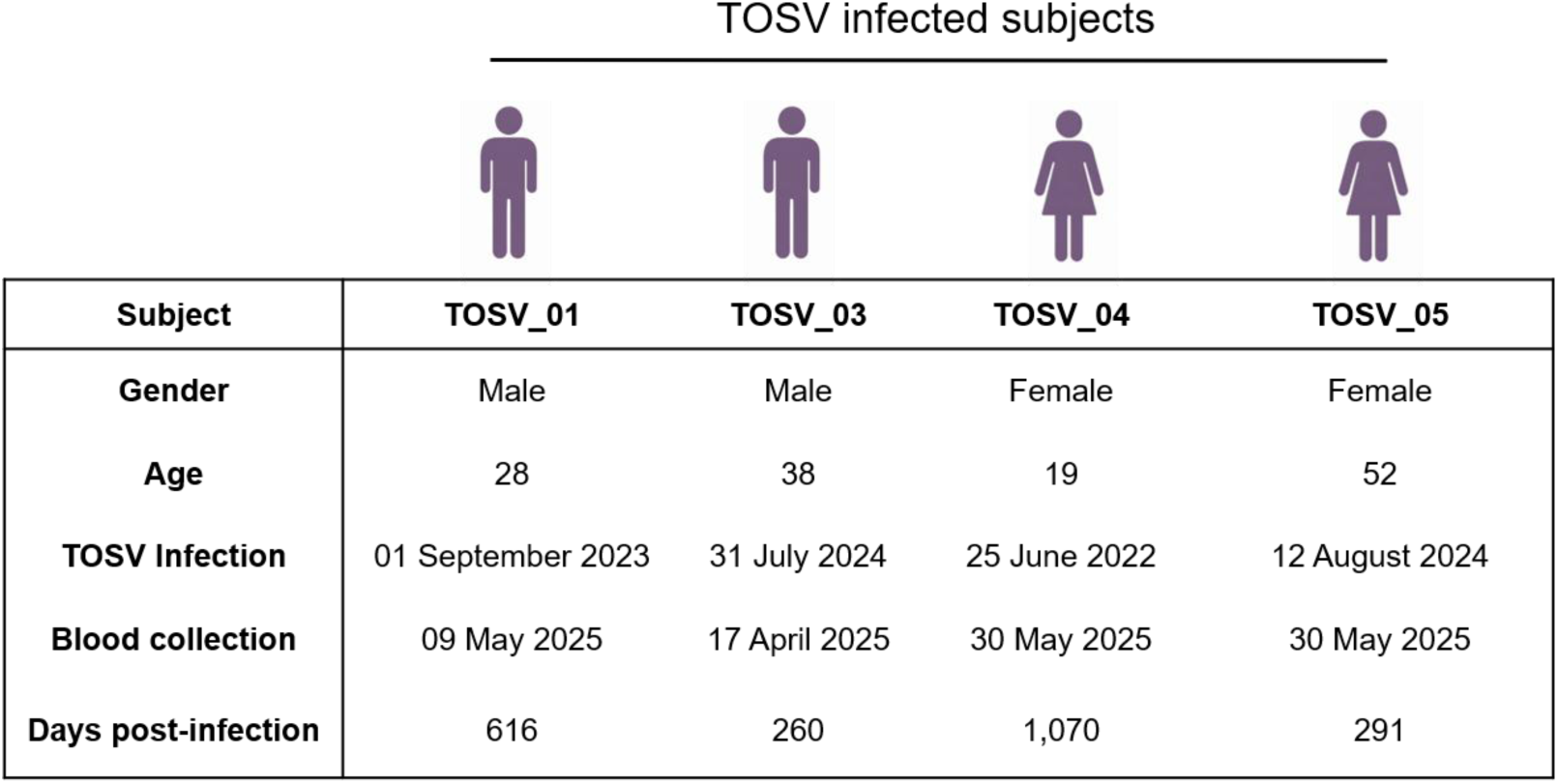
Donor enrollment and sample collection. Schematic overview of donor enrollment and sample collection for TOSV infected individuals (n=4).

**Figure S2.**
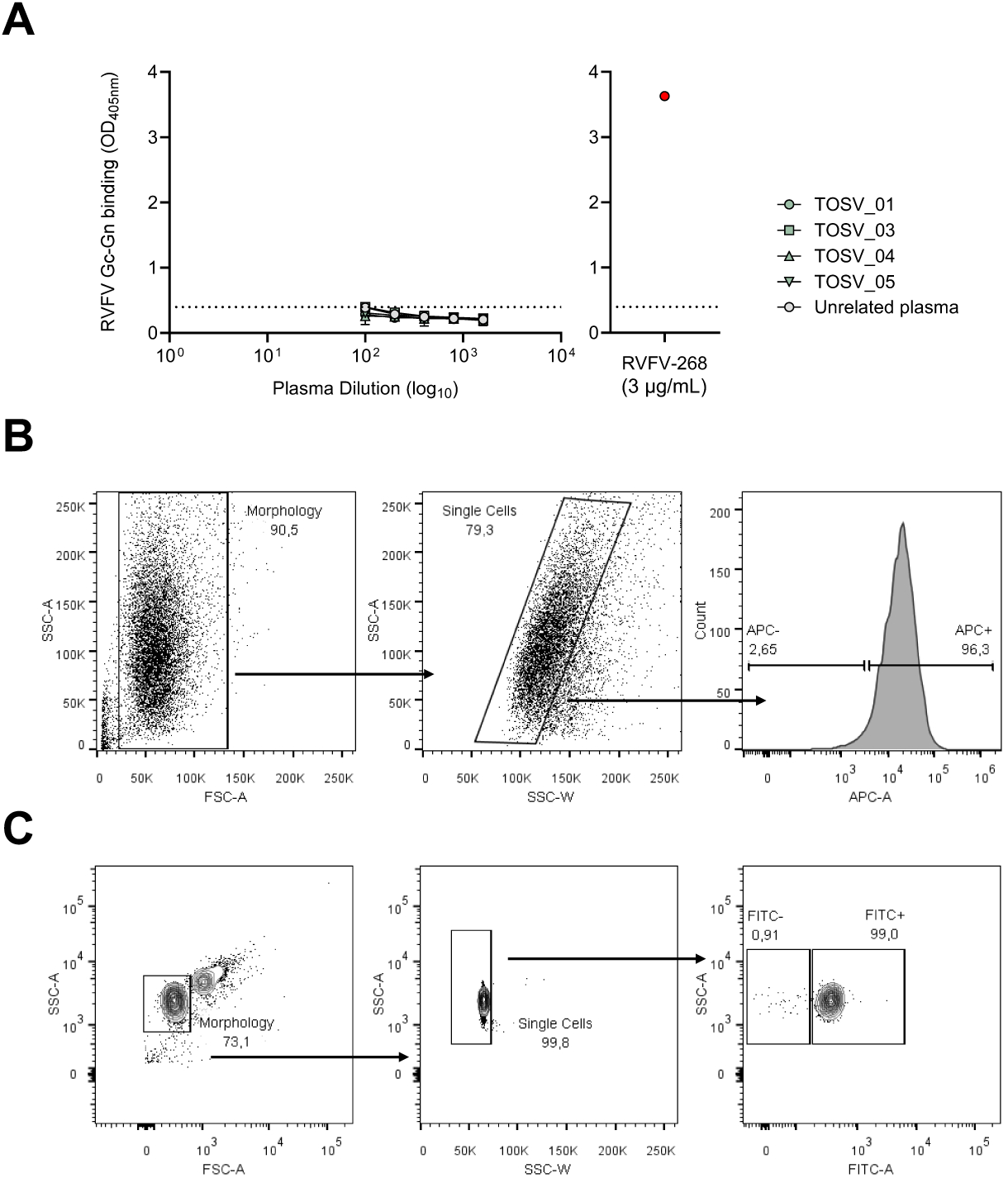
Cross-reactivity of plasma to RVFV and gating strategy for ADCP and ADCD assays. (**A**) Graph showing the ability of plasma samples to bind the RVFV Gn-Gc glycoprotein. Technical replicates were performed and threshold of positivity was set based on the unrelated plasma used as negative control (left panel). RVFV-268 mAb was used as positive control (right panel). (**B**) The gating strategy used to measure the uptake of Gn-Gc-beads by THP-1 cells composed of: morphology, single cells, APC^+^ cells. (**C**) The gating strategy for ADCD analysis in flow cytometry shows morphology, single cells and FITC^+^ subsets cells of positive plasma.

**Figure S3.**
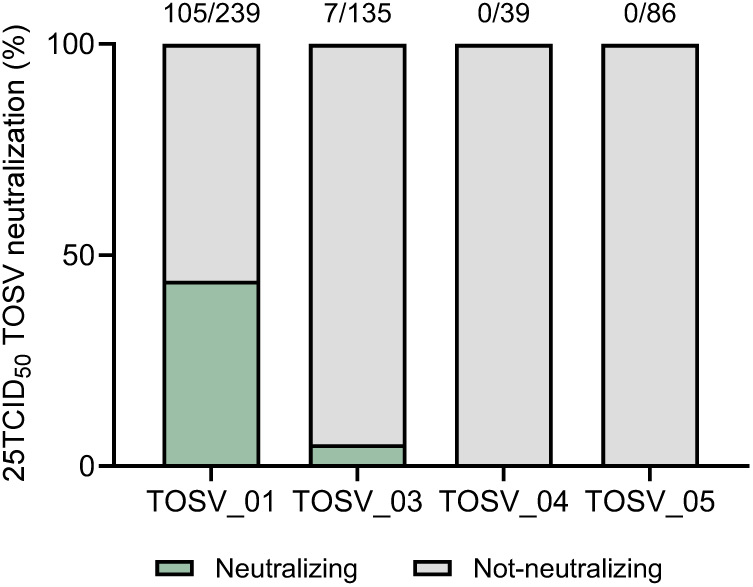
Initial neutralization screening of the 499-expressed mAbs against authentic TOSV (25TCID_50_).

**Table S1.** Details of the 5 antibodies tested *in vivo*.

| Antibody ID | TOSV<br>neutralization -<br>IC100 (ng/mL) | ELISA RVFV<br>Gc-Gn | ELISA RVFV<br>Gc-Gn (OD) | ADCP<br>(MFI) | ADCP<br>(Level) | ADCD<br>(MFI) | ADCD<br>(Level) | heavy chain<br>v_call | heavy chain<br>d_call | heavy chain<br>j_call | light chain<br>v_call | light chain<br>j_call |
| --- | --- | --- | --- | --- | --- | --- | --- | --- | --- | --- | --- | --- |
| R2P2_TOSV_295 | 8,9 | Negative | 0,23026 | 1814831,6 | High | 42330,9 | High | IGHV6-1 | IGHD3-16 | IGHJ6-1 | IGKV3-20 | IGKJ4-1 |
| R2P2_TOSV_311 | 17,7 | Negative | 0,22551 | 529293,6 | Medium | 14773 | High | IGHV3-23 | IGHD3-22 | IGHJ2-1 | IGKV1-33 | IGKJ4-1 |
| R2P2_TOSV_362 | 11,9 | Positive | 0,44666 | 163569,6 | Medium | 24425 | High | IGHV6-1 | IGHD3-16 | IGHJ6-1 | IGKV3-20 | IGKJ4-1 |
| R2P2_TOSV_387 | 253,5 | Positive | 0,78496 | 19521,9 | Low | 13834,8 | High | IGHV4-61 | IGHD2-21 | IGHJ4-1 | IGKV1-5 | IGKJ2-1 |
| R2P2_TOSV_395 | 302,2 | Positive | 1,81421 | 7482,4 | No | 19425,6 | High | IGHV4-31 | IGHD3-10 | IGHJ6-1 | IGKV4-1 | IGKJ2-1 |

## References

Abudurexiti, A., Adkins, S., Alioto, D., Alkhovsky, S. V., Avšič-Županc, T., Ballinger, M. J., Bente, D. A., Beer, M., Bergeron, É., Blair, C. D., Briese, T., Buchmeier, M. J., Burt, F. J., Calisher, C. H., Cháng, C., Charrel, R. N., Choi, I. R., Clegg, J. C. S., de la Torre, J. C.,…Kuhn, J. H. (2019). Taxonomy of the order Bunyavirales: update 2019. Archives of Virology, 164(7), 1949–1965. 10.1007/s00705-019-04253-6

Allen, E. R., Krumm, S. A., Raghwani, J., Halldorsson, S., Elliott, A., Graham, V. A., Koudriakova, E., Harlos, K., Wright, D., Warimwe, G. M., Brennan, B., Huiskonen, J. T., Dowall, S. D., Elliott, R. M., Pybus, O. G., Burton, D. R., Hewson, R., Doores, K. J., & Bowden, T. A. (2018). A Protective Monoclonal Antibody Targets a Site of Vulnerability on the Surface of Rift Valley Fever Virus. Cell Reports, 25(13), p3750–3758.e3754. 10.1016/j.celrep.2018.12.001

Ando, T., Nabeshima, T., Inoue, S., Tun, M. M. N., Obata, M., Hu, W., Shimoda, H., Kurihara, S., Izumikawa, K., Morita, K., Hayasaka, D., Ando, T., Nabeshima, T., Inoue, S., Tun, M. M. N., Obata, M., Hu, W., Shimoda, H., Kurihara, S.,…Hayasaka, D. (2021). Severe Fever with Thrombocytopenia Syndrome in Cats and Its Prevalence among Veterinarian Staff Members in Nagasaki, Japan. Viruses, 13(6), 1142. 10.3390/v13061142

Andreano, E., Nicastri, E., Paciello, I., Pileri, P., Manganaro, N., Piccini, G., Manenti, A., Pantano, E., Kabanova, A., Troisi, M., Vacca, F., Cardamone, D., Santi, C. D., Torres, J. L., Ozorowski, G., Benincasa, L., Jang, H., Genova, C. D., Depau, L.,…Rappuoli, R. (2021). Extremely potent human monoclonal antibodies from COVID-19 convalescent patients. Cell, 184(7), 1821–1835. 10.1016/j.cell.2021.02.035

Andreano, E., Paciello, I., Piccini, G., Manganaro, N., Pileri, P., Hyseni, I., Leonardi, M., Pantano, E., Abbiento, V., Benincasa, L., Giglioli, G., De Santi, C., Fabbiani, M., Rancan, I., Tumbarello, M., Montagnani, F., Sala, C., Montomoli, E., & Rappuoli, R. (2021). Hybrid immunity improves B cells and antibodies against SARS-CoV-2 variants. Nature, 600(7889), 530–535. 10.1038/s41586-021-04117-7

Anywaine, Z., Lule, S. A., Hansen, C., Warimwe, G., & Elliott, A. (2022). Clinical manifestations of Rift Valley fever in humans: Systematic review and meta-analysis. PLOS Neglected Tropical Diseases, 16(3), e0010233. 10.1371/journal.pntd.0010233

Ayhan, N., & Charrel, R. N. (2020). An update on Toscana virus distribution, genetics, medical and diagnostic aspects. Clinical Microbiology and Infection, 26(8), 1017– 1023. 10.1016/j.cmi.2019.12.015

Ayhan, N., Prudhomme, J., Laroche, L., Bañuls, A.-L., Charrel, R. N., Ayhan, N., Prudhomme, J., Laroche, L., Bañuls, A.-L., & Charrel, R. N. (2020). Broader Geographical Distribution of Toscana Virus in the Mediterranean Region Suggests the Existence of Larger Varieties of Sand Fly Vectors. Microorganisms, 8(1), 114. 10.3390/microorganisms8010114

Azerigyik, F. A., Cagle, S. M., Wilson, W. C., Mitzel, D. N., Kading, R. C., Azerigyik, F. A., Cagle, S. M., Wilson, W. C., Mitzel, D. N., & Kading, R. C. (2025). The Temperature-Associated Effects of Rift Valley Fever Virus Infections in Mosquitoes and Climate-Driven Epidemics: A Review. Viruses, 17(2), 217. 10.3390/v17020217

Barash, S., Wang, W., & Shi, Y. (2002). Human secretory signal peptide description by hidden Markov model and generation of a strong artificial signal peptide for secreted protein expression. Biochemical and Biophysical Research Communications, 294(4). 10.1016/S0006-291X(02)00566-1

Braito, A., Corbisiero, R., Corradini, S., Marchi, B., Sancasciani, N., Fiorentini, C., & Ciufolini, M. G. (1997). Evidence of Toscana virus infections without central nervous system involvement: A serological study. European Journal of Epidemiology, 13(7), 761–764. 10.1023/A:1007422103992

Caminade, C., McIntyre, K. M., & Jones, A. E. (2018). Impact of recent and future climate change on vector-borne diseases. Annals of the New York Academy of Sciences, 1436(1), 157–173. 10.1111/nyas.13950

Cartwright, H. N., Barbeau, D. J., & McElroy, A. K. (2021). Isotype-Specific Fc Effector Functions Enhance Antibody-Mediated Rift Valley Fever Virus Protection In Vivo. mSphere, 6(5), 10.1128. 10.1128/msphere.00556-21

Cassetti, M. C., Pierson, T. C., Patterson, L. J., Bok, K., DeRocco, A. J., Deschamps, A. M., Graham, B. S., Erbelding, E. J., & Fauci, A. S. (2023). Prototype Pathogen Approach for Vaccine and Monoclonal Antibody Development: A Critical Component of the NIAID Plan for Pandemic Preparedness. The Journal of infectious diseases, 227(12), 1433–1441. 10.1093/infdis/jiac296

Chapman, N. S., Hulswit, R. J. G., Westover, J. L. B., Stass, R., Paesen, G. C., Binshtein, E., Reidy, J. X., Engdahl, T. B., Handal, L. S., Flores, A., Gowen, B. B., Bowden, T. A., & James E Crowe, J. (2023). Multifunctional human monoclonal antibody combination mediates protection against Rift Valley fever virus at low doses. Nature Communications, 14(1), 5650. 10.1038/s41467-023-41171-3

Chapman, N. S., Zhao, H., Kose, N., Westover, J. B., Kalveram, B., Bombardi, R., Rodriguez, J., Sutton, R., Genualdi, J., LaBeaud, A. D., Mutuku, F. M., Pittman, P. R., Freiberg, A. N., Gowen, B. B., Fremont, D. H., & James E Crowe, J. (2021). Potent neutralization of Rift Valley fever virus by human monoclonal antibodies through fusion inhibition. Proceedings of the National Academy of Sciences of the United States of America, 118(14), e2025642118. 10.1073/pnas.2025642118

Charrel, R. N., Bichaud, L., & Lamballerie, X. d. (2012). Emergence of Toscana virus in the mediterranean area. World Journal of Virology, 1(5), 135. 10.5501/wjv.v1.i5.135

Depaquit, J., Grandadam, M., Fouque, F., Andry, P., & Peyrefitte, C. (2010). Arthropod-borne viruses transmitted by Phlebotomine sandflies in Europe: a review. Eurosurveillance, 15(10). 10.2807/ese.15.10.19507-en

Dessau, M., & Modis, Y. (2013). Crystal structure of glycoprotein C from Rift Valley fever virus. Proceedings of the National Academy of Sciences, 110(5), 1696–1701. 10.1073/pnas.1217780110

DiLillo, D. J., Tan, G. S., Palese, P., & Ravetch, J. V. (2014). Broadly neutralizing hemagglutinin stalk–specific antibodies require FcγR interactions for protection against influenza virus in vivo. Nature Medicine, 20(2), 143–151. 10.1038/nm.3443

Elliott, R. M. (2009). Bunyaviruses and climate change. Clinical Microbiology and Infection, 15(6), 510–517. 10.1111/j.1469-0691.2009.02849.x

Fatma, F., Price, D. A., Rush, R. E., Allen, C., Stuckmeyer, H., Brown, G., Brett, T. J., Hartman, A. L., Leung, D. W., & Amarasinghe, G. K. (2026). Biochemical Basis for LRP1 Interaction With Rift Valley Fever Virus Glycoprotein and Its Role in Viral Entry. Journal of Molecular Biology, 438(19). 10.1016/j.jmb.2026.169907

Gandolfo, C., Prathyumn, S., Terrosi, C., Anichini, G., Savellini, G. G., Corti, D., Bracci, L., Lanzavecchia, A., Roman-Sosa, G., & Cusi, M. G. (2021). Identification of a Neutralizing Epitope on TOSV Gn Glycoprotein. Vaccines, 9(8), 924. 10.3390/vaccines9080924

Giudicelli, V., Chaume, D., & Lefranc, M.-P. (2005). IMGT/GENE-DB: a comprehensive database for human and mouse immunoglobulin and T cell receptor genes. Nucleic Acids Research, 33(suppl_1), D256–D261. 10.1093/nar/gki010

Guardado-Calvo, P., & Rey, F. A. (2017). The Envelope Proteins of the Bunyavirales. Advances in virus research, 98, 83–118. 10.1016/bs.aivir.2017.02.002

Gunn, B. M., Yu, W.-H., Karim, M. M., Brannan, J. M., Herbert, A. S., Wec, A. Z., Halfmann, P. J., Fusco, M. L., Schendel, S. L., Gangavarapu, K., Krause, T., Qiu, X., He, S., Das, J., Suscovich, T. J., Lai, J., Chandran, K., Zeitlin, L., Crowe, J. E.,…Alter, G. (2018). A Role for Fc Function in Therapeutic Monoclonal Antibody-Mediated Protection against Ebola Virus. Cell Host & Microbe, 24(2), 221–233.e225. 10.1016/j.chom.2018.07.009

Hagberg, A. A., Schult, D. A., & Swart, P. J. (2008). Exploring Network Structure, Dynamics, and Function using NetworkX. Python in Science Conference. 10.25080/TCWV9851

Halldorsson, S., Behrens, A.-J., Harlos, K., Huiskonen, J. T., Elliott, R. M., Crispin, M., Brennan, B., & Bowden, T. A. (2016). Structure of a phleboviral envelope glycoprotein reveals a consolidated model of membrane fusion. Proceedings of the National Academy of Sciences, 113(26), 7154–7159. 10.1073/pnas.1603827113

Hao, M., Zhang, G., Zhang, S., Chen, Z., Chi, X., Dong, Y., Fan, P., Liu, Y., Chen, Y., Song, X., Liu, S., Yu, C., Li, J., & Xia, X. (2020). Characterization of Two Neutralizing Antibodies against Rift Valley Fever Virus Gn Protein. Viruses, 12(3), 259. 10.3390/v12030259

Heiden, J. A. V., Yaari, G., Uduman, M., Stern, J. N. H., O’Connor, K. C., Hafler, D. A., Vigneault, F., & Kleinstein, S. H. (2014). pRESTO: a toolkit for processing high-throughput sequencing raw reads of lymphocyte receptor repertoires. Bioinformatics, 30(13), 1930–1932. 10.1093/bioinformatics/btu138

Hornak, K. E., Lanchy, J.-M., & Lodmell, J. S. (2016). RNA Encapsidation and Packaging in the Phleboviruses. Viruses, 8(7), 194. 10.3390/v8070194

Huang, J., Doria-Rose, N. A., Longo, N. S., Laub, L., Lin, C.-L., Turk, E., Kang, B. H., Migueles, S. A., Bailer, R. T., Mascola, J. R., & Connors, M. (2013). Isolation of human monoclonal antibodies from peripheral blood B cells. Nature protocols, 8(10), 1907– 1915. 10.1038/nprot.2013.117

Hulswit, R. J. G., Paesen, G. C., Bowden, T. A., & Shi, X. (2021). Recent Advances in Bunyavirus Glycoprotein Research: Precursor Processing, Receptor Binding and Structure. Viruses, 13(2), 353. 10.3390/v13020353

Kitandwe, P. K., McKay, P. F., Kaleebu, P., & Shattock, R. J. (2022). An Overview of Rift Valley Fever Vaccine Development Strategies. Vaccines, 10(11), 1794. 10.3390/vaccines10111794

Liang, X., Teng, A., Braun, D. M., Felgner, J., Wang, Y., Baker, S. I., Chen, S., Zelphati, O., & Felgner, P. L. (2002). Transcriptionally Active Polymerase Chain Reaction (TAP): high throughput gene expression using genome sequence data. Journal of Biological Chemistry, 277(5), 3593–3598. 10.1074/jbc.M110652200

Liu, Q., He, B., Huang, S.-Y., Wei, F., & Zhu, X.-Q. (2014). Severe fever with thrombocytopenia syndrome, an emerging tick-borne zoonosis. The Lancet Infectious Diseases, 14(8), 763–772. 10.1016/S1473-3099(14)70718-2

Lofano, G., Gorman, M. J., Yousif, A. S., Yu, W.-H., Fox, J. M., Dugast, A.-S., Ackerman, M. E., Suscovich, T. J., Weiner, J., Barouch, D., Streeck, H., Little, S., Smith, D., Richman, D., Lauffenburger, D., Walker, B. D., Diamond, M. S., & Alter, G. (2018). Antigen-specific antibody Fc glycosylation enhances humoral immunity via the recruitment of complement. Science Immunology, 3(26), eaat7796. 10.1126/sciimmunol.aat7796

Mehand, M. S., Al-Shorbaji, F., Millett, P., & Murgue, B. (2018). The WHO R&D Blueprint: 2018 review of emerging infectious diseases requiring urgent research and development efforts. Antiviral Research, 159, 63–67. 10.1016/j.antiviral.2018.09.009

Nicoletti, L., Renzi, A., Caciolli, S., Bartolozzi, D., Balducci, M., Traini, E., Leoncini, F., Baldereschi, M., Paci, P., Padovani, P., Ciufolini, M. G., & Verani, P. (1991). Central nervous system involvement during infection by Phlebovirus toscana of residents in natural foci in central Italy (1977-1988). American Journal of Tropical Medicine and Hygiene, 45(4), 429–434. 10.4269/AJTMH.1991.45.429

Paciello, I., Maccari, G., Pantano, E., Andreano, E., & Rappuoli, R. (2024). High-resolution map of the Fc functions mediated by COVID-19-neutralizing antibodies. Proceedings of the National Academy of Sciences, 121(3), e2314730121. 10.1073/pnas.2314730121

Rolin, A. I., Berrang-Ford, L., & Kulkarni, M. A. (2019). The risk of Rift Valley fever virus introduction and establishment in the United States and European Union. Emerging Microbes & Infections, 2(12), e81. 10.1038/emi.2013.81

Saphire, E. O., Schendel, S. L., Gunn, B. M., Milligan, J. C., & Alter, G. (2018). Antibody-mediated protection against Ebola virus. Nature Immunology, 19(11), 1169–1178. 10.1038/s41590-018-0233-9

Savellini, G. G., Genova, G. D., Terrosi, C., Bonito, P. D., Giorgi, C., Valentini, M., Docquier, J.-D., & Cusi, M. G. (2008). Immunization with Toscana virus N-Gc proteins protects mice against virus challenge. Virology, 375(2), 521–528. 10.1016/j.virol.2008.02.006

Schreur, P. J. W., Kormelink, R., & Kortekaas, J. (2018). Genome packaging of the Bunyavirales. Current opinion in virology, 33, 151–155. 10.1016/j.coviro.2018.08.011

Seo, J.-W., Kim, D., Yun, N., & Kim, D.-M. (2021). Clinical Update of Severe Fever with Thrombocytopenia Syndrome. Viruses, 13(7), 1213. 10.3390/v13071213

Spiegel, M., Plegge, T., & Pöhlmann, S. (2016). The Role of Phlebovirus Glycoproteins in Viral Entry, Assembly and Release. Viruses, 8(7), 202. 10.3390/v8070202

Sui, L., Sun, P., Liu, N., Zhao, Y., Zhao, Y., Zheng, J., & Liu, Q. (2025). The public health threat of emerging phenuiviruses. One Health, 20, 101055. 10.1016/j.onehlt.2025.101055

Tiller, T., Meffre, E., Yurasov, S., Tsuiji, M., Nussenzweig, M. C., & Wardemann, H. (2008). Efficient generation of monoclonal antibodies from single human B cells by single cell RT-PCR and expression vector cloning. Journal of immunological methods, 329(1-2), 112–124. 10.1016/j.jim.2007.09.017

Turk, Y. K., Ergunay, K., Kohl, A., Hughes, J., & McKimmie, C. S. (2024). Toscana virus – an emerging Mediterranean arbovirus transmitted by sand flies. Journal of General Virology, 105(11), 002045. 10.1099/jgv.0.002045

Verani, P., Ciufolini, M. G., Nicoletti, L., Balducci, M., Sabatinelli, G., Coluzzi, M., Paci, P., & Amaducci, L. (1982). Ecological and epidemiological studies of the Toscana virus, an arbovirus isolated from phlebotomines. Ann Ist Super Sanita, 18(3), 397–399.

Wang, Q., Ma, T., Wu, Y., Chen, Z., Zeng, H., Tong, Z., Gao, F., Qi, J., Zhao, Z., Chai, Y., Yang, H., Wong, G., Bi, Y., Wu, L., Shi, R., Yang, M., Song, J., Jiang, H., An, Z.,…Yan, J. (2019). Neutralization mechanism of human monoclonal antibodies against Rift Valley fever virus. Nature Microbiology, 4(7), 1231–1241. 10.1038/s41564-019-0411-z

Yamanaka, A., Kirino, Y., Fujimoto, S., Ueda, N., Himeji, D., Miura, M., Sudaryatma, P. E., Sato, Y., Tanaka, H., Mekata, H., & Okabayashi, T. (2020). Direct Transmission of Severe Fever with Thrombocytopenia Syndrome Virus from Domestic Cat to Veterinary Personnel. Emerging infectious diseases, 26(12), 2994–2998. 10.3201/eid2612.191513

Ye, J., Ma, N., Madden, T. L., & Ostell, J. M. (2013). IgBLAST: an immunoglobulin variable domain sequence analysis tool. Nucleic Acids Research, 41(W1), W34–W40. 10.1093/nar/gkt382

Yoshikawa, T. (2021). Vaccine Development for Severe Fever with Thrombocytopenia Syndrome. Viruses, 13(4), 627. 10.3390/v13040627

Zhao, C., Hao, M., Bian, T., Zhao, X., Chi, X., Chen, Z., Fu, G., Zhu, Z., Fang, T., Yu, C., Li, J., & Chen, W. (2025). Screening of Neutralizing Antibodies Targeting Gc Protein of RVFV. Viruses, 17(4), 559. 10.3390/v17040559

